# The role of conscious attention in statistical learning: evidence from patients with impaired consciousness

**DOI:** 10.1101/2024.01.08.574591

**Authors:** Lucas Benjamin, Di Zang, Ana Fló, Zengxin Qi, Pengpeng Su, Wenya Zhou, Liping Wang, Xuehai Wu, Peng Gui, Ghislaine Dehaene-Lambertz

**Author notes:** Corresponding authors: Ghislaine Dehaene-Lambertz and Peng Gui. These authors contributed equally to this work (Lucas Benjamin, Di Zang).

## Abstract

The debate over whether conscious attention is necessary for statistical learning has produced mixed and conflicting results. Testing individuals with impaired consciousness may provide some insight, but very few studies have been conducted due to the difficulties associated with testing such patients. In this study, we examined the ability of patients with varying levels of consciousness disorders (DOC), including coma, unresponsive wakefulness syndrome, minimally conscious patients, and emergence from minimally conscious state patients, to extract statistical regularities from an artificial language composed of four randomly concatenated pseudowords. We used a methodology based on frequency tagging in EEG, which was developed in our previous studies on speech segmentation in sleeping neonates. Our study had two main objectives: firstly, to assess the automaticity of the segmentation process and explore correlations between the level of covert consciousness and the abilities to extract statistical regularities, second, to explore a potential new diagnostic indicator to aid in patient management by examining the correlation between successful statistical learning markers and consciousness level. We observed that segmentation abilities were preserved in some minimally conscious patients, suggesting that statistical learning is an inherently automatic low-level process. Due to significant inter-individual variability, word segmentation may not be a sufficiently robust candidate for clinical use, unlike temporal accuracy of auditory syllable responses, which correlates strongly with coma severity. Therefore, we propose that frequency tagging of an auditory stimulus train, a simple and robust measure, should be further investigated as a possible metric candidate for DOC diagnosis.

## Introduction

The analysis of the statistical patterns present in sensory sequences, a mechanism called statistical learning, enables us to uncover the structure of the input and to build an efficient mental model of the environment. Such a mechanism operating between adjacent syllables, can be exploited to detect words in a continuous linguistic stream ***(Saffran et al., 1996)***. In this type of behavioral experiment, four pseudo words are randomly concatenated to form a continuous sequence with high predictability of the next syllable within the pseudo word but a drop of transition probability between pseudo-words. Despite the absence of other cues for word segmentation, such as those provided by prosodic indices, participants subsequently distinguished words composed of syllables with high transition probabilities, from part-words comprising a low transition between syllables. This result has been extended to non-linguistic streams ***(Saffran et al., 1999; Schön et al., 2008)*** as well as to visual sequences ***(Fiser and Aslin, 2002)***. It is neither specific to the human species ***(Hauser et al., 2001; James et al., 2020)***, nor limited to adjacent elements, extending to non-adjacent dependencies ***(Kabdebon et al., 2015)***. It has also been shown that humans can learn higher-order statistical structures in sequences ***(Benjamin et al., 2023a; Dehaene et al., 2015; Garvert et al., 2017; Schapiro et al., 2013)***. Thus, statistical learning has been proposed to be a fundamental learning mechanism operating in different modalities and sensitive to statistical regularities at different temporal scales.

Whether statistical learning occurs automatically and whether this mechanism requires attention remains a matter of debate, making it unclear under what circumstances such learning can occur. While most studies have used passive exposure to sequences in awake and attentive subjects, the impact of attention on statistical learning has been explicitly questioned by some authors, with mixed results. While some studies have shown a drastic decrease in performance under divided attention ***(Fernandes et al., 2010; Toro et al., 2005)***, others have shown preserved segmentation of auditory sequence when attention was diverted by asking subjects to focus on an independent visual sequence ***(Batterink and Paller, 2019; Benjamin et al., 2021)***. Another study even reports improved performances in participants under cognitive fatigue after performing an effortful working memory task prior to the sequence presentation (Smalle et al., 2022). Thus, studies that directly manipulate attentional focus do not provide any definitive answer regarding the interaction between statistical learning and participants’ attentional resources.

Another angle of attack of this question is to study statistical learning in sleeping subjects. Although not directly testing attention, sleep studies address the automaticity of this mechanism. Batterink et al (2022) recently tested sleeping adults with pseudo-word segmentation tasks composed of either bi-syllabic or tri-syllabic pseudo-words. They reported that sleeping adults were only sensitive to bi-syllabic pairs but failed to extract tri-syllabic words. It suggests that sleep might reduce the integration period of the statistical computation without preventing the computation between adjacent items. This interpretation appears congruent with Strauss and colleagues’ study (2015) who observed during sleep, a preserved mismatch response to local auditory violations of a sequence regularity but no reaction to a violation concerning a regularity at a longer time scale, that induced a P300 in awake adults. In contrast, one-to three-day-old neonates tested with EEG in a pseudo-word segmentation paradigm similar to that of Batterink et al in adults ***(Benjamin et al., 2022; Fló et al., 2019, 2022a)*** not only calculated statistics between syllables, but also segmented tri-syllabic pseudowords during sleep (for quadrisyllabic pseudo-words see ***(Benjamin et al., 2023b)***).

A more rigorous test of the automaticity of statistical learning would be to examine this learning in comatose patients. Among Disorders of Consciousness (DOC), patients may exhibit varying degrees of residual consciousness, as assessed by diagnostic tools such as CRS-R ***(Giacino et al., 2004)*** or categorical classification into Unresponsive Wakefulness Syndrome (UWS) -no sign of visible awareness- or Minimally Conscious State (MCS) –visible partial awareness-***(Bruno et al., 2011; Giacino et al., 2002)***. One attempt to measure statistical learning in DOC patients was recently made by Xu et al **(2022)** using bi-syllabic words concatenated in a continuous, monotonic stream. They reported some learning in patients with emerging consciousness. Indeed, the power at the frequency of syllable pairs and its harmonics were significantly above zero, suggesting successful segmentation of sequences into word pairs. Although this study supports the possibility of statistical learning in DOC patients, it had some limitations. First, the bi-syllabic units used in the study, which required only a pairing between two items, do not encompass all aspects of segmentation, as shown in the sleep studies discussed above. Furthermore, the pairs presented in the study could be either frequent and meaningful in natural language or reversed and thus meaningless (e.g., “go home” versus “home go”). However, even in the reversed condition, syllables within a pair are more related to each other in natural language than syllables between pairs. This makes it difficult to differentiate between current learning of statistical regularities in the stream and a reactivation in memory of previous exposure to natural language. This limitation was partially addressed by obtaining similar results with the learning of artificial tri-syllabic pseudowords, but the sample of minimally conscious patients was small (N=8).

In the present study, we used the experimental design and techniques of our previous research in sleeping neonates ***(Fló et al., 2022a)*** (see Methods): Our aim was to investigate the ability of DOC patients to recognize statistical regularities in an artificial syllable sequence composed of four tri-syllabic pseudo-words that were concatenated in a pseudo-random manner (fig1). The syllables within the words followed each other predictively (transitional probabilities = 1), while between the words, the transitional probabilities dropped to 1/3. In addition, the subjects were also presented with a random sequence composed of the same syllables but without forming pseudo-words, which had a flat transitional probability of 1/11. We also included a group of healthy awake adults as a control to compare with the clinical population.

**Figure 1:**
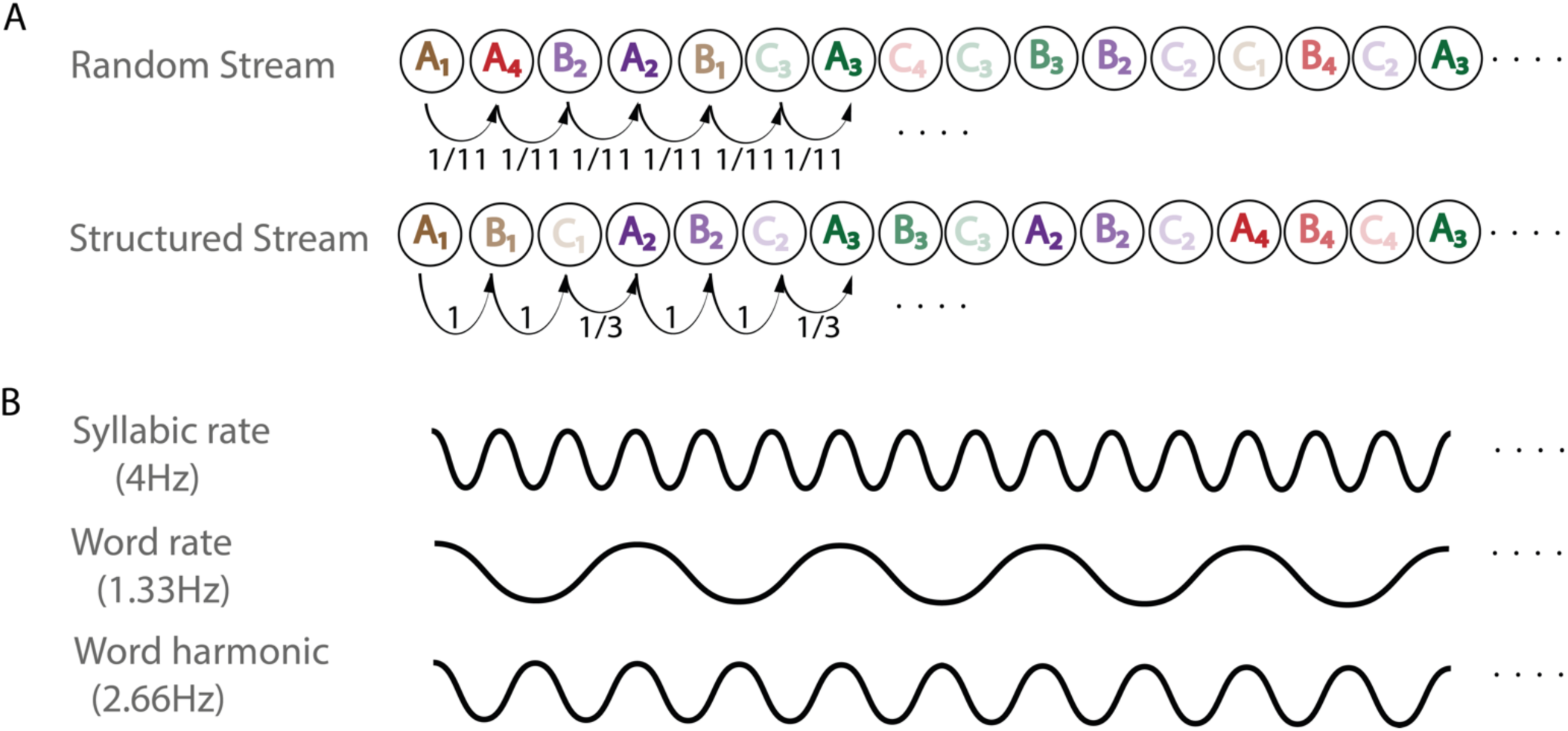
Presentation of the paradigm and dataset. (A) Description of the two streams presented to participants. The random stream consists of syllables that can be followed by any of the other 11 syllables, resulting in a flat transitional probability of 1/11 throughout the sequence. The structured stream is composed of four tri-syllabic pseudo-words with transition probabilities between syllables equal to 1 inside the pseudo-words and 1/3 between the pseudo-words. (B). Frequency tagging analyses : The syllables were presented at a rate of 4Hz, which is expected to elicit a 4Hz oscillation in the brain of normal hearing subjects. If, and only if, the structured sequence was segmented based on transition probabilities, the phase locking value (PLV) at the word rate (1.33Hz) and its harmonic (2.66Hz) should increase relative to the random stream.

We utilized high-density EEG recordings (256 channels) to measure sequence segmentation with the frequency tagging method, as it has been demonstrated to be a robust approach for evaluating non-responsive subjects in our previous studies ***(Fló et al., 2022a)***. This technique relies on detecting rhythmicity of the brain activity driven by the rhythmicity of the input sequence. It permits the monitoring of sequence segmentation by following the modulation of power and Phase Locking Value (PLV) at the syllabic and word frequencies. Analyzing the power at the syllabic rate allowed us to assess basic auditory processing at the individual level and check which patients have intact auditory perception. Segmentation of the tri-syllabic words is revealed by an increase in the word-rate frequency and eventually harmonics.

Not only does this paradigm address the question of conscious attention in statistical learning, but also offers a promising way to enhance the clinical assessment of covert consciousness in DOC patients. Previous studies have found modification of EEG features with the level of consciousness, such as spectral power ***(Goldfine et al., 2011)***, auditory ERP amplitude and latency ***(Strauss et al., 2015)***, and mismatch responses in EEG oddball paradigms ***(Laureys and Schiff, 2012)***. The depth of language processing during time-locked natural language exposure has also been suggested as a useful metric for predicting patient outcomes ***(Gui et al., 2020)***. Given the importance of language function ***(Claassen et al., 2019; Gui et al., 2020; Sanz et al., 2021; Sokoliuk et al., 2021)*** and correct metabolic functioning of the left middle temporal cortex ***(Aubinet et al., 2020)*** in the recovery of DOC patients, investigating language related paradigms is a promising venue for new clinical tools. With the present study, we also aim to investigate whether neural markers of statistical learning can be utilized for diagnosis and outcome prediction in DOC patients.

## Methods

### Participants

81 patients from Huashan hospital with disorders of consciousness were included in the experiment for a total of 180 recordings from March 2021 to September 2022. Clinical assessment of the patient state was made just before each recording. Out of those 180 recordings, 14 were classified as coma, 65 as UWS and 99 as minimally conscious or emerging of consciousness. 4 recordings were discarded as participants were labeled as fully recovered. We then rejected recordings that had not at least one good epoch in each condition (random and structured): 3 recordings were rejected based on this criterion leaving a total of 172 recordings. 26 healthy control adults (>18yo, average age:25.27; 18 females and 9 males) from local community were also tested along which 24 completed the experiments (72 recordings). Similar to the patient group, we rejected 3 recordings with no good epoch in one condition, and thus analyzed the 69 remaining recordings. The experiment consisted of three stimulus sequences (see details in the following), of which 78 patients completed 88 recordings of list 1, 50 patients completed list 2 recordings and 46 patients completed list 3 recordings. Therefore, a total of 40 patients completed three different lists at different time points. In the healthy control group, 24 healthy volunteers completed all three lists.

### Stimuli

We synthesized the syllables using the open-source text-to-speech synthesizer eSpeaker ***(Jonathan and Reece, 2020)*** with Mandarin Chinese language (zh) and the female voice variant f2. We set the pitch parameter to 70 and adapted the speed to obtain syllables with a duration as close as possible to 250 ms. We further corrected the syllables using Praat ***(Boersma and Weenink, 2020)***. Specifically, we removed silent periods in the beginning and the end to obtain syllables lasting exactly 250ms and removed pitch changes setting a constant pitch of 225 Hz. The syllables audio files were concatenated without pauses to obtain the streams, and the first and last 4.5 s were ramped up and down to avoid the start and end of the stream might serve as perceptual anchors.

The structured streams consisted of a semi-random concatenation of four tri-syllabic pseudowords. The only restriction for concatenation was that the same pseudoword could not appear twice in a row, and the same two pseudowords could not repeatedly alternate more than two times (i.e., the sequence WkWjWkWj, where Wk and Wj are two words, was forbidden). We balanced phonetic features across the three syllables of the pseudowords to avoid they could serve as segmentation cues. In addition, we created three different structured streams by changing the arrangement of the 12 syllables (each syllable occupied a different position within the pseudoword in each stream). The random stream resulted from concatenating the 12 syllables semi-randomly (without syllables repetition), giving an average uniform TP of 1/11. We also created test words to be presented in isolation but we do not present the results here.

### Procedure

Scalp electrophysiological activity was recorded using a 256-electrodes net (GTEN 200, Magstim EGI) referred to the vertex with a sampling frequency of 1000 Hz. The recording procedure was similar to the one used in neonates ***(Fló et al., 2022a)***. Participants first heard a random sequence of 4 minutes (960 syllables) followed by a structured stream of 4 minutes (960 syllables, 320 words). After, participants listen to 8 repetitions of 30s (120 syllables, 40 words) of structured sequences followed by a block of 16 pseudo-words test trials. Patients were told to pay attention to the auditory stimuli.

All Healthy subjects were tested with the three structured streams (lists) on different days. For the DOC patients, we tried to follow the same pattern by testing each subject on each list as much as the hospital constraints allowed us. On average DOC subjects were tested 2.27 times, with 40 subjects having been tested on the three lists. We obtained 88 recordings with list 1, 50 with list 2 and 46 with list 3.

### Data preprocessing

Data were resampled to 250 Hz, band-pass filter 0.3–30 Hz and pre-processed using APICE pipeline for MATLAB ***(Fló et al., 2022b)*** with default rejection strength parameters. This preprocessing pipeline detects artifacts on the continuous signal based on deviance from the distribution of data points. It later corrects isolated artifacts with interpolation methods, recovering part of the signal. Long periods of artifacts, or simultaneous failure of several channels cannot be corrected with neighbors’ data and the corresponding epochs are removed from further analysis.

In order to remove eye movements and blinks, we also performed ICA on the healthy control data.

### Frequency tagging

The pre-processed data were segmented from the beginning of each sequence into segments comprising 13 words to approach 10s long epochs (13*0.750=9.75s). Segments were not overlapping to avoid an artefact in the frequency domain related to the length of the overlap ***(Benjamin et al., 2021)***. Epochs with artifacts were rejected. Data were converted to the frequency domain using the Fast Fourier Transform (FFT) algorithm and the Phase Locking Value (PLV) and Power were estimated for each electrode in both random and structured conditions. The phase locking value ranges from 0 (completely asynchronized data) to 1 (completely timed-locked activity). The value at each frequency bin estimation was then normalized by subtracting the mean value of eight neighboring frequency bins on each side.

### Statistical analyses

They were performed on the three frequencies of interest: 4Hz corresponding to the syllabic rate, 1.33 and 2.66 corresponding to the frequency of the trisyllabic words and its harmonic. Results of the analyses performed on the PLV are presented in the main text, and for power in the supplementary material. Results are largely similar for these two metrics.

For each electrode and for each frequency, it was first tested whether the PLV (and power) was above 0 with a one-sample t-test against zero, second whether these values for the word frequencies were larger during the structured stream relative to the random stream (structured > random one-way paired t-test). In all analyses, p-values were corrected for multiple comparisons (256 electrodes) using FDR (fig 2).

**Figure 2:**
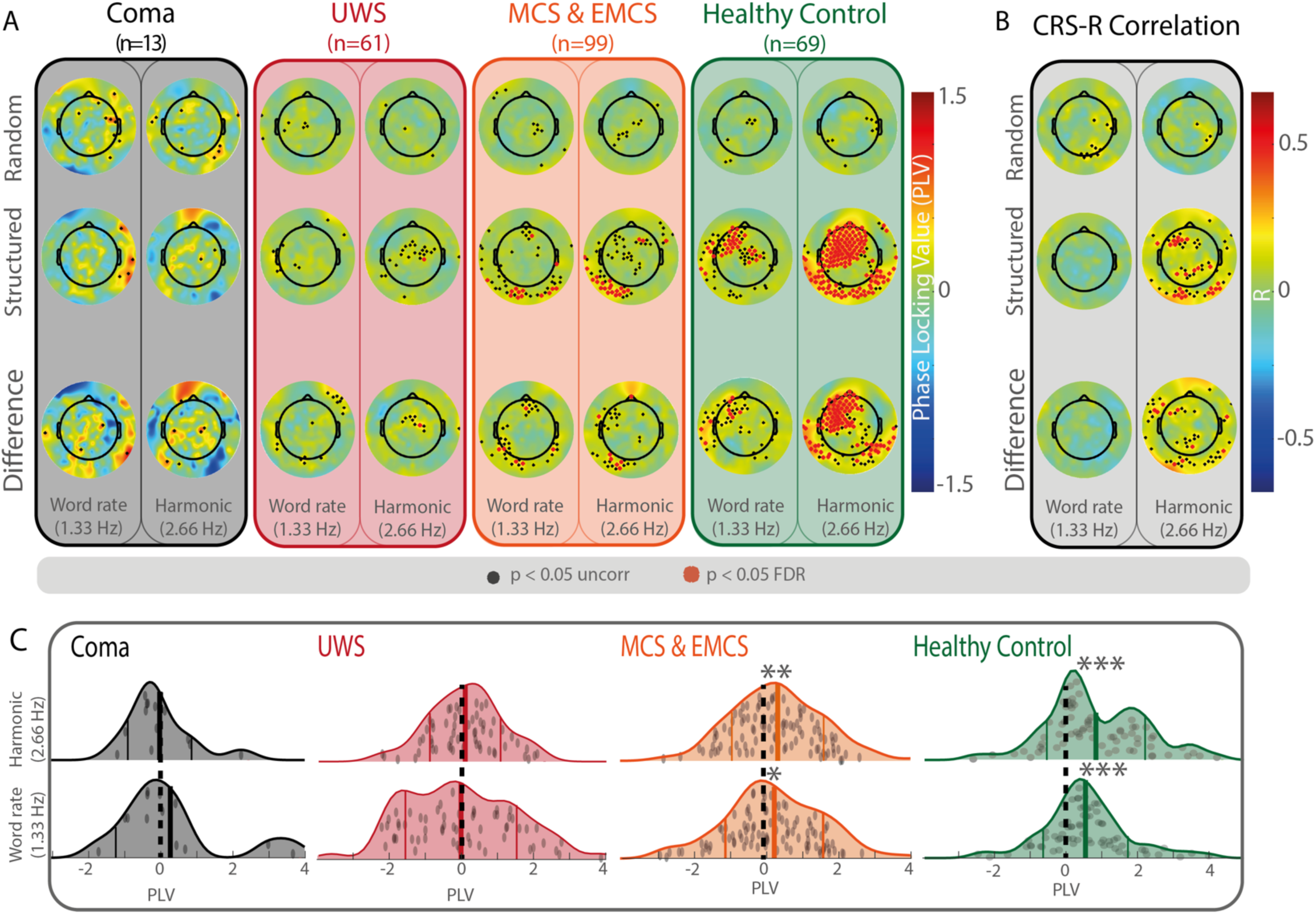
Neural entrainment analysis. (A) Normalized Phase Locking Value (PLV) for each electrode at the word rate (1.33Hz) and at its first harmonic (2.66Hz) during the random and structured streams in the recordings of Coma, UWS, MCS and Healthy subjects. The bottom line presents the topography of the contrast structured > random. Dots represent electrodes with p< 0.05 before multiple comparison correction and red crosses electrodes significant after FDR correction. (B) Correlation of the PLV at word rate and its harmonic with Comatose Recovery Scale Revised (CRS-R). The harmonic of the word rate (2.66Hz) significantly correlates with the clinical score only during the structured stream. (C) Distribution of the effect size for each recording in each group and for the word rate and its harmonics. Each dot represents the difference between structured and random PLV at the word and the harmonic rates for the 10 best electrodes in each condition of one particular recording. Significant difference of the distribution average with zero is represented by stars (* <p<0.05, ** p<0.01, ***p< 0.001).

### Individual effect size

We then estimated the effect size for each recording to determine whether we could distinguish patients who performed the task from those who did not segment the structured sequence. To do this, we calculated the difference at the frequency of interest between the 10 electrodes with the highest PLV in the structured condition and the 10 electrodes with the highest PLV in the random condition. The reason for selecting the 10 best electrodes per recording was to obtain a robust measurement of the effect by selecting the most effective electrodes without being affected by possible differences in topography at the individual level induced by different brain lesions.

We performed this analysis at the word rate (1.33Hz) and its first harmonic (2.66 Hz) and used one-way tests to compare the average of the distribution with zero (no PLV difference between structured and random sequences).

### Modulation of the neural responses by level of consciousness

To examine how the responses were influenced by the level of consciousness in DOC patients, we computed correlations between the Phase Locking Value (PLV) of each electrode and the Coma Recovery Scale-Revised (CRS-R) score measured prior to the recording in DOC patients only. Importantly, healthy controls were not included in this analysis to ensure that any observed correlation was not solely driven by the differences between healthy and DOC subjects, but rather by the varying degrees of consciousness within the DOC group.

### Modulation of words segmentation by the syllabic response

To be able to segment words in such a stream, patient should at least have a minimal auditory function and some patients may not meet this criterion. We considered the responses at the syllabic rate as a proxy of low-level auditory function. Therefore, the previous analyses were done again retaining only patients with a positive average syllabic rate PLV, that is patients for whom the mean PLV across all electrodes for both the random and structured conditions was > 0 (fig 3). This rejection metric is quite stringent as it rejected participants with impaired auditory processing but also recording with too low signal/noise ratio to correctly detect this metric. We also re-calculated the correlation between the PLV at the word rates and level of consciousness after regressing out the PLV at the syllabic rate for each electrode: we first performed a regression between word rate PLV and the syllabic rate PLV for each electrode; then the residuals of these regressions were correlated with CRS-R (fig3 panel D).

**Figure 3:**
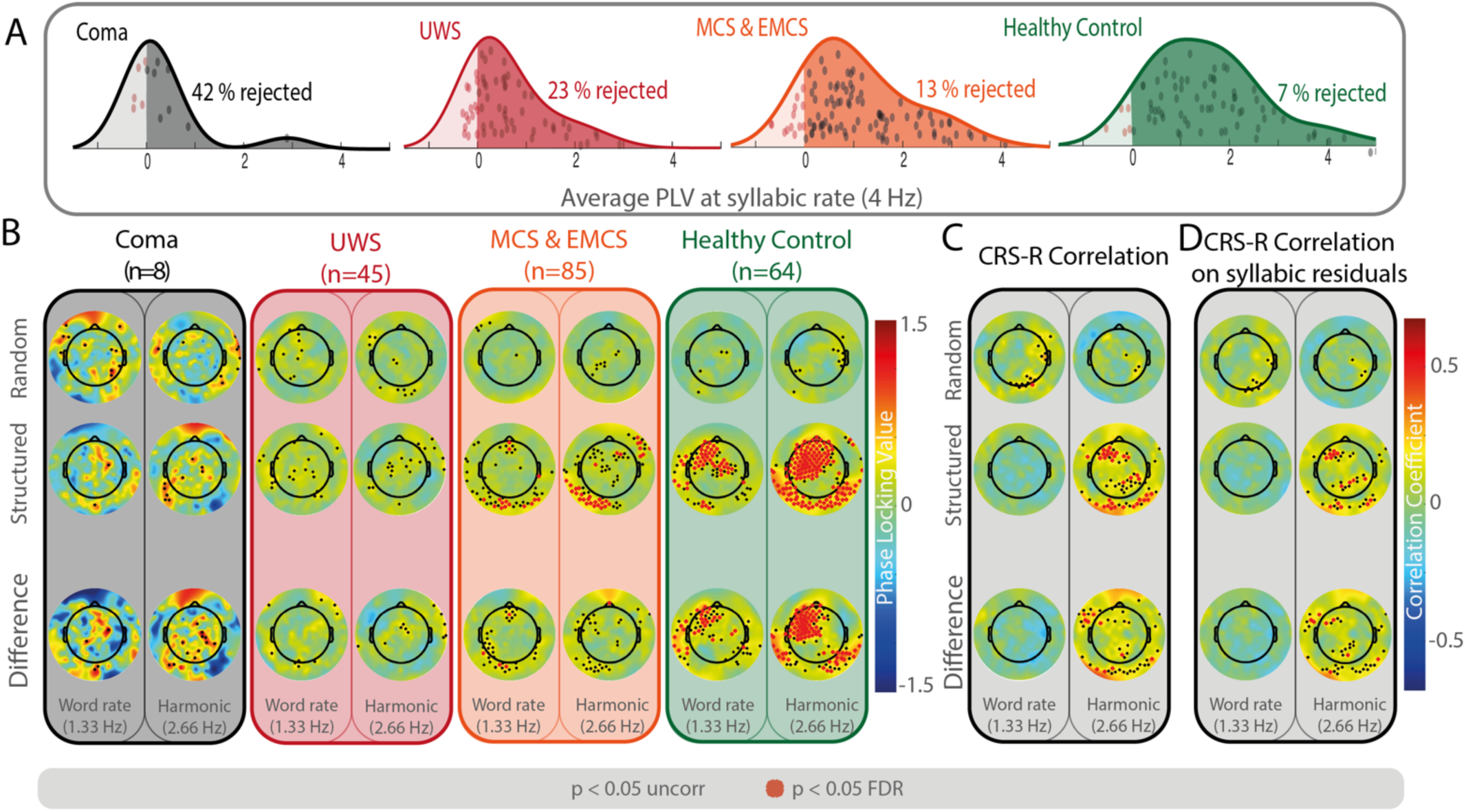
(A) Average PLV at the syllabic rate for each recording. Syllabic rate is a good metric of preserved auditory perception and correct signal/noise ratio in the recording. For the following analysis, we only kept the recordings with a positive average syllabic rate as we cannot be sure that the other participants even heard the stimuli. Red dots represent recording with negative syllabic rates, who were then excluded from the analyses presented in B and C. (B) Normalized PLV at each electrode at the word rate (1.33Hz) and its first harmonic (2.66Hz) during the random and structured streams in patients with Coma, UWS, MCS and Healthy subjects, excluding recordings with negative average PLV at syllabic rate (red dots on Panel A). The bottom line is the contrast structured > random. Dots represent electrodes with p< 0.05 before multiple comparison correction and red crosses electrodes significant after FDR correction. (C) Correlation of the PLV at word rate and harmonic with CRS-R scores excluding recordings with negative average PLV at syllabic rate. The harmonic of the word rate (2.66Hz) significantly correlates with this score during the structured stream only. (D) To account for the variation of neural entrainment at syllabic rate, we computed the correlation after having regressed out the effect of DOC on syllable entrainment, results remained similar

## Results

### 1) Evidence of word segmentation in the different experimental groups

To estimate whether patients were able to correctly segment the words concatenated in the structured stream, we measured the normalized PLV at the word rate and its first harmonic and compared the values to those measured in the random stream (fig 2 panel A).

First, in healthy control subjects, we found many electrodes with significant positive PLV at both word rate and its harmonic when they listened to the structured stream (p < 0.05 FDR corrected) but not to the random stream, with a significant difference between the two conditions in many electrodes (Figure 2). The same analysis in patients in the minimally conscious state (clinical assessment: MCS or EMCS) showed similar results, although weaker in terms of number of significant electrodes. In the UWS group, a trend in the same directions as the other groups was visible (Structured stream at 2.66Hz: 23 channels with p<0.05 uncorrected), but only one channel survived the FDR correction at the word rate harmonic. The comparison with the random stream showed a very modest effect (1 electrode with p < 0.05 FDR corrected). In the Coma patients, no electrode showed a significant segmentation effect.

### 2) Correlation of the segmentation performance with CRS-R

We then estimated for each electrode the correlation of the word and harmonic PLV with the clinical Coma recovery scale revised (CRS-R) estimated just before the recording, excluding the healthy participants. We found a highly significant correlation spread on many channels of the neural entrainment (normalized PLV) and the CRS-R score during the structured stream only for the word first harmonic (24 channels with p < 0.05 FDR) with a significant difference with the random condition (figure 2B).

### 3) Individual effect size

We estimated the effect size for each recording to assess inter-individual variability and to determine whether we could separate subjects who segmented the structured sequence from those who did not (fig2C). This analysis replicated our previous results, with both healthy controls and MCS patients showing an average effect size greater than zero for both the word rate and its harmonic. The obtained distribution showed a high inter-individual variability with no clear bi-modal distribution thus not enabling to confidently conclude on patients’ ability to perform the task at the individual level.

### 4) Controls taking into account low-level auditory perception as captured by the response at the syllabic rate

The previous correlation between PLV at the word harmonics and CRS-R score might indicate either a modulation in statistical learning and segmentation abilities with DOC severity or a spurious correlation due to a higher number of patients with auditory perception impairment and a severe DOC. Fortunately, PLV at the syllabic rate effectively summarizes basic auditory perception and temporal synchronization of auditory ERP. We presented the distribution of the average PLV at syllabic rate for each recording in figure 3A. Therefore, we repeated the previous analyses (fig 2) while excluding recordings with negative syllabic rates to ensure that all recordings left are with patients with correct hearing and a level of signal/noise in the recording that enable correct measure of frequency tagging. The results remained similar, both considering the PLV measures in each group (fig3 panel B) and the correlation with CRS-R (fig3 panel C).

Finally, to mitigate the impact of the variation of the PLV at the syllabic rate of each electrode on the segmentation metrics, we regressed out the syllabic rate for each recording for each electrode before computing the correlation of the residuals with CRS-R score. For this analysis, we included all recordings of DOC patients, comprising those with negative average syllabic rates. We obtained results similar to the original analyses: 1) a significant correlation during the structured stream only with the first harmonic, but not at the fundamental; 2) a significant contrast between structured and random sequence streams (fig3 panel D).

The analyses presented in fig 2 and 3 have been replicated using power instead of PLV with similar results (fig S1 and S2 in supplementary material).

### 5) Auditory ERP measured with syllabic rate neural entrainment

Independently from statistical learning, we found that the neural entrainment at the syllabic rate was highly correlated with the clinical assessment of the level of consciousness in DOC patients. Indeed, we found many electrodes for which PLV and Power at the syllabic rate (4hz) robustly correlated with diagnostic scales, such as CRS-R (fig 4). PLV at the syllabic rate significantly correlated with CRS-R on most electrodes. On fig 4 B, we display the distribution of the average PLV value from the significant electrodes, for each subject by clinical assessment group. We clearly observe higher PLV at the syllabic rate associated with better clinical assessment. We also compared for each group the probability of improved outcome six months later depending on the PLV at the syllabic, word and word harmonic rates and found no significant differences (for the syllabic rate analysis, recordings associated with an improved six-month outcome are displayed in green on fig4).

**Figure 4:**
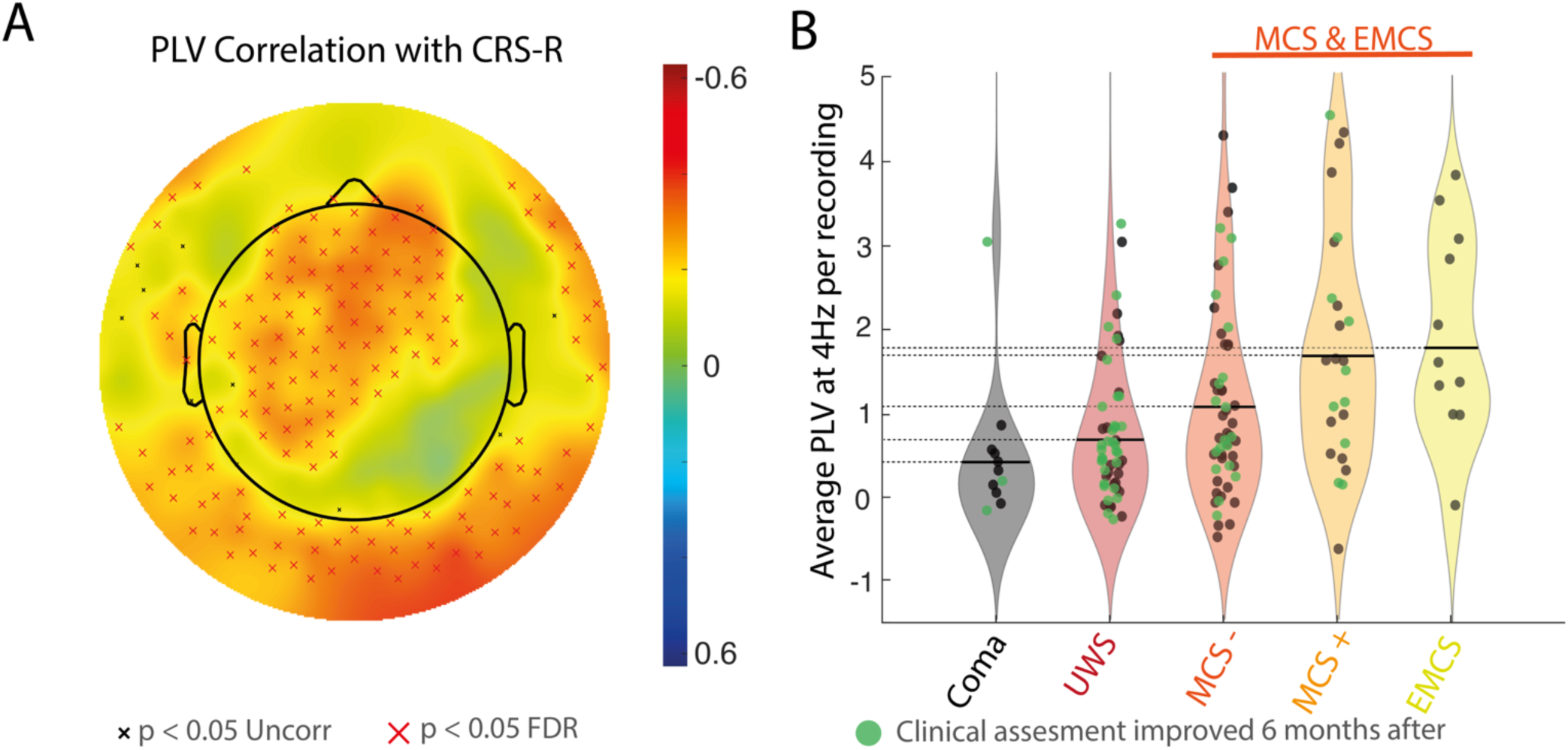
Syllabic rate correlation with CRS-R. (A) For each electrode, we correlated the PLV measure at the syllabic rate (average across random and structured condition) with patients’ CRS-R measured just before the recording. (B) For better visualization of the effect, we report here the distribution of the average PLV at 4Hz across electrodes for each recording and patients’ status. The average neural entrainment is modulated by participant’s coma depth but the inter-recording variance within each diagnostic group stays higher than the variance explained by the coma depth. Note that this figure is not independent from the analysis done in Panel A and so we did not perform statistical analysis on this. This is just presented for a better visualization and estimation of the inter-recording variance. Recordings associated with an improved clinical assessment 6 months later are displayed in green. We found not systematic relation between PLV at the syllabic rate and probability of improved clinical condition 6 months later.

## Discussion

Frequency tagging provides a direct neural measure for monitoring word segmentation within a continuous speech stream obviating the need of an explicit behavioral response. This approach has proven its usefulness in assessing segmentation abilities in preverbal infants ***(Kabdebon et al., 2015)*** and neonates ***(Fló et al., 2022a)***. Consequently, we proposed to apply this paradigm in DOC patients as also did Xu et al (2022). Our first goal was to establish whether statistical learning and auditory sequence segmentation might persist in patients with disorder of consciousness amidst conflicting literature on the role of conscious attention. Our second goal was to explore if statistical learning metrics may help clinical diagnosis and care of these patients.

### Statistical learning is partially preserved in DOC patients

The automaticity of statistical learning is still being debated. Indeed, while some studies showed a large decline in performance under divided attention ***(Fernandes et al., 2010; Toro et al., 2005)*** arguing for the need of focused attention on the task, others have reported that sequence segmentation persists even when outside the focus of attention ***(Batterink and Choi, 2021; Batterink and Paller, 2019; Benjamin et al., 2021)*** and even improves with cognitive fatigue that impairs focal attention (Smalle et al., 2022). Furthermore, sleeping neonates can automatically segment a stream based on its statistical properties ***(Benjamin et al., 2023b; Fló et al., 2022a)***. Similarly, other studies investigating visual statistical learning with hypnosis ***(Nemeth et al., 2013)***, or TMS disruption of DLPFC ***(Ambrus et al., 2020)***, suggest that statistical learning abilities are enhanced when prefrontal activity is reduced.

Studies in DOC patients provide valuable insight on this issue. In a recent study, ***(Xu et al., 2022)*** showed that some comatose patients were able to extract bi-syllabic real words (or mixed) arguing for a preserved minimal version of statistical learning on pairs of real words. Here, we extend and strengthen this result by using a stream of tri-syllabic pseudo-words with flat intonation, i.e. without any additional linguistic cues to aid segmentation. Thus, any increase of PLV and power at the frequency of three syllables can only be based on the online calculation of the statistical relationship, or transitional probability (TP), between the syllables, i.e. TP=1 between syllables belonging to the same word and TP=1/3 between syllables belonging to different words. We checked for any spurious effects by comparing this structured stream with a stream consisting of a concatenation of the same syllables with a flat probability (TP=1/11). Significant differences between the two streams were observed in MCS & EMCS patient group and possibly a very weak trend in UWS patients. Since MCS patients suffer from severe attentional dysfunction, this result provides evidence that the full focus of attention is not needed for this type of learning. Sleeping adults were only able to chunk bisyllabic pseudowords unlike neonates who succeeded with trisyllabic pseudo-words as is the case here. It can be related to the linguistic expert adults’ bias to segment the stream in shorter units (Franck et al, 2010) that might interfere more in a sleep stage than in coma. In any case, chunking a stream in its tri-syllabic components reveals that even MCS & EMC patients were able to integrate TP over several syllables to discover the TP drop that is used to chunk words.

### Why are results more visible on the first harmonic compared to the word rate?

All our analyses pointed toward a greater neural entrainment at the first harmonic compared to the fundamental frequency at the word rate. In our design, the harmonic of the word rate (2.66Hz) is different from half of the frequency of the syllabic rate (2Hz); thus, the modulation seen here can only be due to the discovery of the word structure and not to the perception of the syllables. The evoked activity by each syllable and word superposes as there is no pause between syllables and words. The shape of this event-related activity is complex, and the Fourier transform decomposes it into a set of sinusoids with different power depending on the ERP shape. A significant response at the first harmonic argues for a rhythmic response that vanishes faster than the word length, such as a larger response words’ first syllable. In contrast, an activity drooling over the following word (e.g. integration of the three syllables) would be more visible at the fundamental frequency ***(Zhou et al., 2016)***. ERP shapes can explain the different sensitivity of the two measures observed in the above analyses. Further experiments are needed to investigate whether this difference reveals a different encoding of the word in memory. For instance, sleeping neonates segment a tri-syllabic non-word stream but only remember the first syllable of the words, contrary to adults who memorize the entire word ***(Fló et al., 2022a)***.

### Correlation between the level of residual consciousness and statistical learning

We observed a significant correlation between CRS-R score and the first harmonic of the word rate in DOC patients. This reveals that even though this statistical learning task does not require attention, a deeper consciousness disorder penalizes learning. Therefore, we tried to separate whether impaired learning was related to deficient statistical computations or to a deficit in auditory perception due to degraded auditory/phonetic encoding or the result of suboptimal synchronization of cortical activity with the stimulus. To do this, we used the 4Hz entrainment as a metric for the auditory low-level processing quality as well as a proxy of the recording quality which is sometimes impaired by the electrically noisy environment of the hospital wards. We linearly regressed it to CRS-R score. We then used the residuals of this regression in the correlation analysis with the word rate and its harmonic (fig3). Despite this stringent control, a significant correlation remained between CRS-R and the first harmonic only for the structured stream as well as a significant difference between the structured and random condition. This suggests that statistical learning abilities were affected by the degree of residual consciousness, even in cases where the brain exhibited the capacity to track the syllabic rhythm. It remains possible that in some cases, syllabic entrainment was based solely on the vocalic nucleus of the CV syllable, without the exact phonemes being encoded, thus ruling out the possibility of statistical learning. This could be the case for patients with lesions of the left perisylvian regions involved in phonetic processing. Fama et al (2022) reported that stroke patients, notably those with anterior lesions, did not show evidence of statistical learning in a behavioral paradigm in which they had to rate their familiarity with words, part-words and non-words after 10 min of familiarization with a similar structured stream than here. Despite the high number of recordings studied here, we had not enough power to orthogonalize the DOC degree and the type, size, localization of the lesions. Further studies are needed, notably by testing musical tones to simplify the identification of the tokens in the stream.

#### Is there a clinical interest for this type of paradigm?

Despite the significant result described above, a word-segmentation task as implemented here might not be usable as a standalone clinical tool, although it could be relevant to include it in a battery of tests as this task targets a basic learning mechanism. The search for better indices of recovery, as well as indices quantifying the integrity of different brain functions beyond the anatomical lesions visible on MRI, is a necessity to guide care. However, here, the CRS-R effect size observed was smaller than the inter-individual variance and not highly significant and language functions might be better quantified by sentences in the native language ***(Edlow and Naccache, 2021; Gui et al., 2020; Sokoliuk et al., 2021)***. By contrast, the neural entrainment at the syllabic rate proved much more informative. Indeed, the correlations with CRS-R were greatly and highly significant and features of the auditory ERPs have been shown to be usable ***(Strauss et al., 2015)***. In our data-set, many electrodes showed a significant correlation between the auditory 4Hz neural entrainment and CRS-R measures. Steady-State measure is a more robust and time-economic way to elicit brain responses than isolated ERP, and neural entrainment is a more robust way, to look at the same neural response. This is confirmed by the typical auditory topography of the effect size of the correlation (fig 4 A). Previous studies suggested that the brain responses to violation of auditory regularities, such as mismatch negativity or P300 waves ***(Chennu and Bekinschtein, 2012; Qin et al., 2008; Rivera-Lillo et al., 2018; Sanz et al., 2021; Wang et al., 2017)***, can indicate the presence or absence of awareness in these patients. Thus, syllabic rate entrainment is a promising venue. Further research might be useful to better characterize which frequencies to be entrained are the most sensitive and which electrodes are the most informative for clinical application.

## Conclusion

In this study, we discovered evidence of preserved statistical learning of word boundaries in some DOC patients, using neural entrainment measurements. This confirms our hypothesis that attention is not required for statistical learning and extends the previous findings of efficient statistical learning abilities in sleeping neonates. Our study shows that statistical learning is an automatic process that scans the auditory environment even in conditions of disturbed conscious attention. In addition, we showed that these metrics of statistical learning were significantly correlated with diagnostic metrics such as CRS-R, implying that they can be used as indicators of the level of consciousness and the prognosis of DOC patients. Finally, we proposed that neural entrainment robustness could be of interest for better characterization of auditory ERP modification in DOC patients, warranting further investigation of this measure in relation to the neural mechanisms and the clinical markers of consciousness disorders.

## Acknowledgments

We thank all patients and families who accepted to participate in the study. We also would like to thank the Huashan and Hebin hospitals, and the MEG Facility of CAS CEBSIT/ION in data collection. LB thanks the Treilles foundation for its support.

## Supplementary material

In the main text, we presented all analyses with PLV as a measure of neural entrainment. We replicated here all analyses with Power as a metric for neural entrainment estimation. The results are mainly similar to those presented in the main text with PLV.

Figure S1 corresponds to fig 2 and presents power analyses. The results were similar with those presented o, the main text with PLV.

**Figure S1:**
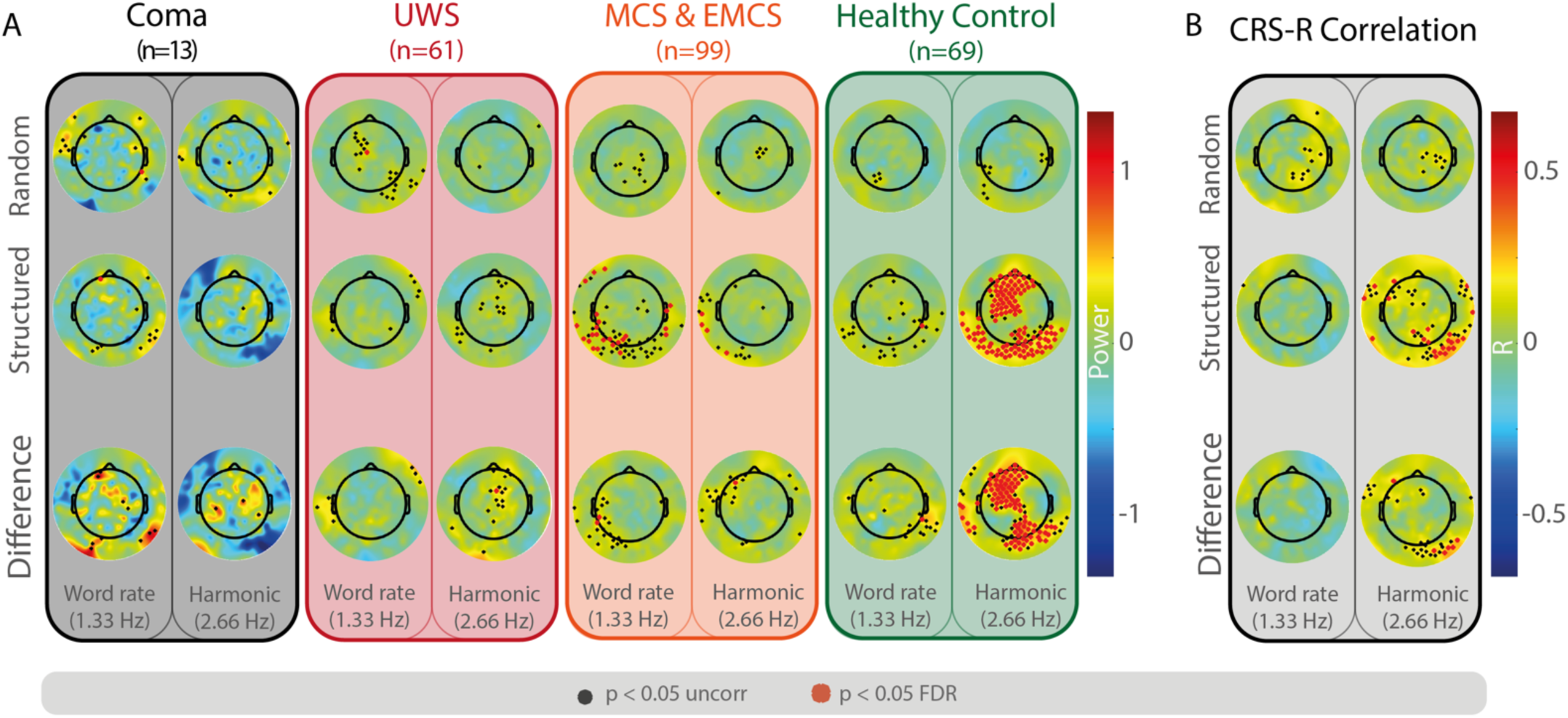
Replication of results from fig 1 with power instead of PVL. A) Power of each electrode at the word rate (1.33Hz) and its first harmonic (2.66Hz) during the random and structured streams. The bottom line present the contrast structured > random. Dot represent electrode with p< 0.05 before multiple comparison correction. The red crosses indicate electrodes significant after FDR correction. (B) Correlation of the Power at word rate and harmonic with Comatose Recovery Scale Revised (CRS-R). The harmonic of the word rate (2.66Hz) significantly correlates with this clinical assessment during the structured stream only.

Figure S2 corresponds to fig 3 and presents power analyses. The results were similar with those presented in the main text with PLV.

**Figure S2:**
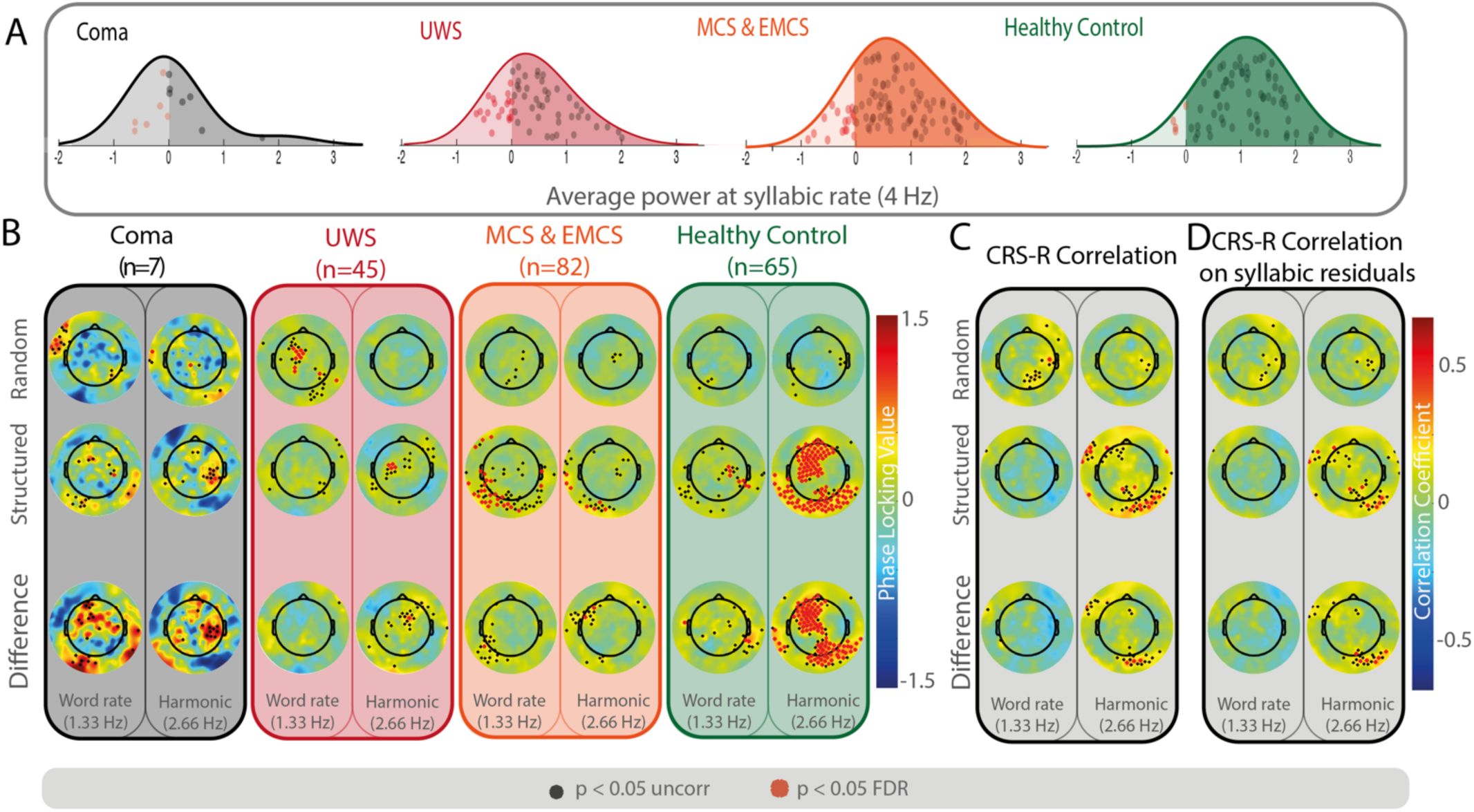
(A) Average syllabic rate Power measure for each recording. Syllabic rate is a good metric for preserved auditory perception. For the following analysis, we only kept the recordings with positive average syllabic rate as we cannot be sure that the other participants even heard the stimuli. Red dots represents recording with negative syllabic rate and were then excluded from further analysis. (B) Power of each electrode at the word rate (1.33Hz) and its first harmonic (2.66Hz) during the random and structured streams for recording of UWs, MCS or Healthy subjects restricted to recordings with positive average Power at syllabic rate (see Panel A). The bottom line is the contrast structured > random to accesses possible increase of Power during the structured compared to the random stream sequences. Dot represent electrode with p< 0.05 before multiple comparison correction. The red crosses indicate electrodes significant after FDR correction. (C) Correlation of the Power at word rate and harmonic with Comatose Recovery Scale Revised (CRS-R) restricted to recordings with positive average Power at syllabic rate (see Panel A). The harmonic of the word rate (2.66Hz) significantly correlates with this clinical assessment during the structured stream only. (D) To account for the variation of syllabic rate, we first computed a regression between word rate (resp harmonic) and the syllabic rate Power per electrode. The residuals of this correlation are then used to correlate with CRS-R. The results are very comparable to the original correlation without syllabic rate regressed out.

